# An atlas of the aging lung mapped by single cell transcriptomics and deep tissue proteomics

**DOI:** 10.1101/351353

**Authors:** Ilias Angelidis, Lukas M. Simon, Isis E. Fernandez, Maximilian Strunz, Christoph H. Mayr, Flavia R. Greiffo, George Tsitsiridis, Elisabeth Graf, Tim-Matthias Strom, Oliver Eickelberg, Matthias Mann, Fabian J. Theis, Herbert B. Schiller

**Affiliations:** Helmholtz Zentrum München, Institute of Lung Biology and Disease, Member of the German Center for Lung Research (DZL), Munich, Germany; Helmholtz Zentrum München, Institute of Computational Biology, Munich, Germany; Helmholtz Zentrum München, Institute of Human Genetics, Munich, Germany; University of Colorado, Department of Medicine, Division of Respiratory Sciences and Critical Care Medicine, Denver, CO, USA; Max Planck Institute of Biochemistry, Department of Proteomics and Signal Transduction, Martinsried, Germany; Department of Mathematics, Technische Universität München, Munich, Germany

**Keywords:** aging, chronic lung disease, single cell transcriptomics, proteomics, extracellular matrix, lipid metabolism, lung cell atlas

## Abstract

Aging promotes lung function decline and susceptibility to chronic lung diseases, which are the third leading cause of death worldwide. We used single cell transcriptomics and mass spectrometry to quantify changes in cellular activity states of 30 cell types and the tissue proteome from lungs of young and old mice. Aging led to increased transcriptional noise, indicating deregulated epigenetic control. We observed highly distinct effects of aging on cell type level, uncovering increased cholesterol biosynthesis in type-2 pneumocytes and lipofibroblasts as a novel hallmark of lung aging. Proteomic profiling revealed extracellular matrix remodeling in old mice, including increased collagen IV and XVI and decreased Fraser syndrome complex proteins and Collagen XIV. Computational integration of the aging proteome and single cell transcriptomes predicted the cellular source of regulated proteins and created a first unbiased reference of the aging lung. The lung aging atlas can be accessed via an interactive user-friendly webtool at: https://theislab.github.io/LungAgingAtlas

---

The intricate structure of the lung enables gas exchange between inhaled air and circulating blood. As the organ with the largest surface area (~70 m^2^ in humans), the lung is constantly exposed to a plethora of environmental insults. A range of protection mechanisms are in place, including a highly specialized set of lung resident innate and adaptive immune cells that fight off infection, as well as several stem and progenitor cell populations that provide the lung with a remarkable regenerative capacity upon injury^1^. These protection mechanisms seem to deteriorate with advanced age, since aging is the main risk factor for developing chronic lung diseases, including chronic obstructive pulmonary disease (COPD), lung cancer, and interstitial lung disease (ILD)^2, 3^. Advanced age causes a progressive impairment of lung function even in otherwise healthy individuals, featuring structural and immunological alterations that affect gas exchange and susceptibility to disease^4^. Aging decreases ciliary beat frequency in mice, thereby decreasing mucociliary clearance and partially explaining the predisposition of the elderly to pneumonia5. Senescence of the immune system in the elderly has been linked to a phenomenon called ‵inflammaging′, which refers to elevated levels of tissue and circulating pro-inflammatory cytokines in the absence of an immunological threat^6^. Several previous studies analyzing the effect of aging on pulmonary immunity point to age dependent changes of the immune repertoire as well as activity and recruitment of immune cells upon infection and injury^4^. Vulnerability to oxidative stress, pathological nitric oxide signaling, and deficient recruitment of endothelial stem cell precursors have been described for the aged pulmonary vasculature^7^. The ECM of old lungs features changes in tensile strength and elasticity, which were discussed to be a possible consequence of fibroblast senescence^8^. Using atomic force microscopy, age-related increases in stiffness of parenchymal and vessel compartments were demonstrated recently^9^, however, the causal molecular changes underlying these effects are unknown.

Aging is a multifactorial process that leads to these molecular and cellular changes in a complicated series of events. The hallmarks of aging encompass cell-intrinsic effects, such as genomic instability, telomere attrition, epigenetic alterations, loss of proteostasis, deregulated nutrient sensing, mitochondrial dysfunction and senescence, as well as cell-extrinsic effects, such as altered intercellular communication and extracellular matrix remodeling^2, 3^. The lung contains at least 40 distinct cell types^10^, and specific effects of age on cell type level have never been systematically analyzed. In this study, we build on rapid progress in single cell transcriptomics^11, 12^, which recently enabled the generation of a first cell type resolved census of murine lungs^13^, serving as a starting point for investigating the lung in distinct biological conditions as shown for lung aging in the present work. We computationally integrated single cell signatures of aging with state of the art mass spectrometry driven proteomics^14^ to generate the first multi-omics whole organ resource of aging associated molecular and cellular alterations in the lung.

## Results

### A single cell atlas of mouse lung aging reveals common deregulated transcriptional control

To generate a cell-type resolved map of lung aging we performed highly parallel genome-wide expression profiling of individual cells using the Dropseq workflow^15^, which uses both molecule and cell-specific barcoding, enabling great cost efficiency and accurate quantification of transcripts without amplification bias^16^. Single cell suspensions of whole lungs were generated from 3 (n=8) and 24 (n=7) month old mice. After quality control, a total of 14,813 cells were used for downstream analysis (Fig. 1a). Unsupervised clustering analysis revealed 36 distinct clusters corresponding to 30 cell types, including all major known epithelial, mesenchymal, and leukocyte lineages (Fig. 1b and d). We observed very good overlap across mouse samples and most clusters were derived from >70% of the mice of both age groups (Fig. S1a). The definition of cell types (clusters in tSNE map) was very comparable between old and young mice, indicating that the cell type identity was not strongly confounded by the aging effects (Fig. S1b). Two clusters exclusively contained cells from a single mouse and were removed from downstream analysis. Interestingly, we identified even rare (<1%, 43 cells) cell types such as megakaryocytes, which were recently identified as a novel unexpected tissue resident cell type in mouse lung^17^. Of note, some samples contributed as little as a single cell to this megakaryocyte cluster, emphasizing the power and accuracy of the computational workflow used here for data integration from multiple mice.

**Figure 1.**
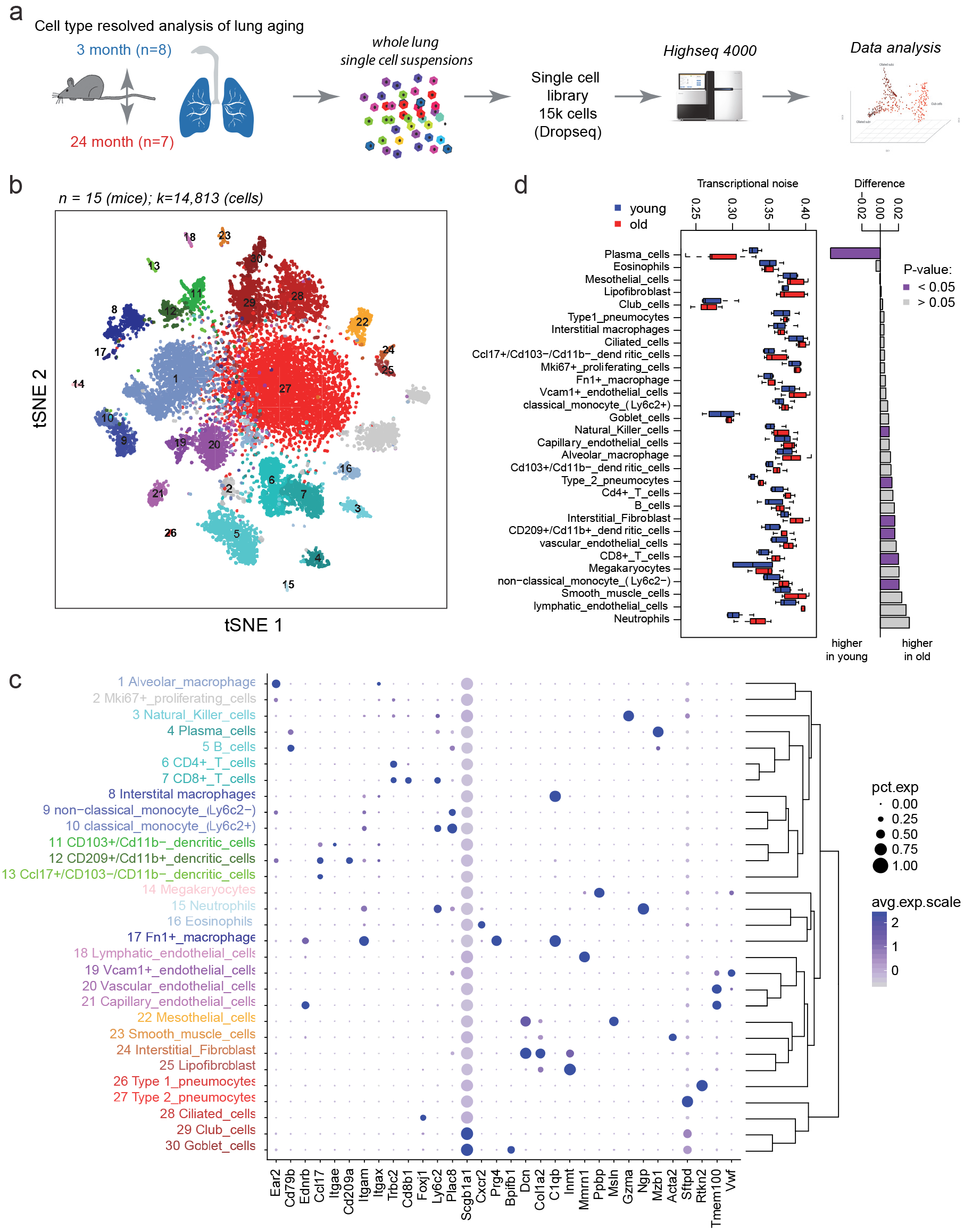
A single cell atlas of mouse lung aging. (a) Experimental design – whole lung single cell suspensions of young and old mice were analyzed using the Dropseq workflow. (b) The tSNE visualization shows unsupervised transcriptome clustering, revealing 30 distinct cellular identities. (c) The dotplot shows (1) the percentage of cells expressing the respective selected marker gene using dot size and (2) the average expression level of that gene based on UMI counts. Rows represent hierarchically clustered cell types, demonstrating similarities of transcriptional profiles. (d) Boxplot illustrates transcriptional noise by age and celltype. Barplot to the right depicts the difference between the median transcriptional noise values between young and old mice. Purple color indicates differences trending towards statistical significant (P: <0.05).

We used differential gene expression analysis to determine cell type specific marker genes with highly different levels between clusters (Fig. 1c, Table S1). The clusters were annotated with assumed cell type identities based on (1) known marker genes derived from expert annotation in literature, and (2) enrichment analysis using Fisher’s exact test of gene expression signatures of isolated cell types from databases including ImmGen^18^ and xCell^19^. Correlation analysis of marker gene signatures revealed that similar cell types clustered together, implying correct cell type annotation (Fig. 1c). We used the matchSCore tool to compare the cluster identities of our dataset with the lung data in the recently published Mouse Cell Atlas^13^, and found very good agreement in cluster identities and annotations (Fig. S1c).

It was suggested that aging is the result of an increase in transcriptional instability rather than a coordinated transcriptional program, and that an aging associated increase in transcriptional noise can lead to ‵fate drifts′ and ambiguous cell type identities^20, 21^. Therefore, we quantified transcriptional noise by calculating the median euclidean distance between the gene expression profile of all single cells from each mouse and the average gene expression profile of the respective cell type. A significant positive association between transcriptional noise and age was observed after accounting for cell type identity as a covariate (Analysis of variance, P: <1e-5). Thus, this analysis confirmed an increase in transcriptional noise with aging in most cell types (Fig. 1d) in line with previous reports in the human pancreas^21^ or mouse CD4+ T cells^20^.

### Multi-omics data integration of mRNA and protein aging signatures

To capture age dependent alterations in both mRNA and protein content, a second independent cohort of young and old mice was analyzed with a state of the art shotgun proteomics workflow (Fig. S2 and Table S2). To compare proteome and transcriptome data we generated *in silico* bulk samples from the scRNA-seq data by summing expression counts from all cells for each mouse individually (Fig. 2a). Differential gene expression analysis of 21969 genes from *in silico* bulks revealed a total of 2362 differentially expressed genes (FDR < 10%) between the two age groups (Fig. 2b, Table S3). From whole lung tissue proteomes we quantified 5212 proteins across conditions and found 213 proteins to be significantly regulated with age, including 32 ECM proteins (FDR < 10%, Fig. 2c, Supplementary Fig. S2 and Table S2).

**Figure 2.**
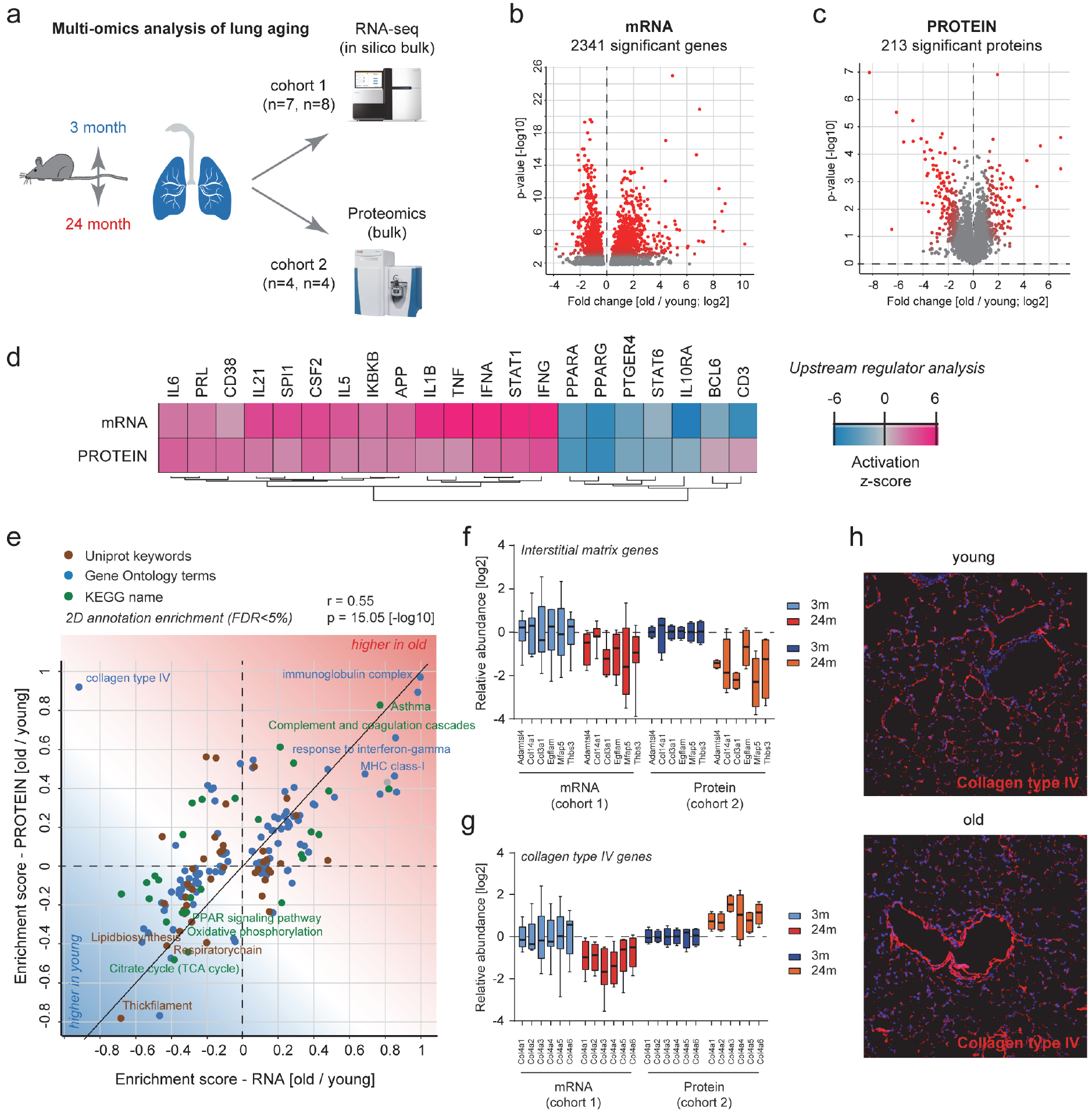
Multi-omics analysis of lung aging. (a) Experimental design-two independent cohorts of young and old mice were analyzed by RNA-seq and mass spectrometry driven proteomics respectively. (b, c) The volcano plots show significantly regulated genes in the (b) transcriptome and (c) proteome. (d) Gene expression and protein abundance fold changes were used to predict upstream regulators that are known to drive gene expression responses similar to the ones experimentally observed. Upstream regulators could be cytokines or transcription factors. The color coded activation z-score illustrates the prediction of increased or decreased activity upon ageing. (e) The scatter plot shows the result of a two dimensional annotation enrichment analysis based on fold changes in the transcriptome (x-axis) and proteome (y-axis), which resulted in a significant positive correlation of both datasets. Types of databases used for gene annotation are color coded as depicted in the legend. (f, g) Normalized relative abundance of the depicted (f) interstitial matrix and (g) Collagen IV genes in the transcriptome and proteome experiments respectively. (h) Immunofluorescence image of collagen type IV using confocal microscopy at 25x magnification. Note, the increased fluorescence intensity around airways in old mice.

Prediction of the upstream regulators^22^ of the observed expression changes in either the transcriptome or proteome data gave very similar results (Fig. 2d). In both datasets from independent mouse cohorts, we discovered a pro-inflammatory signature, which included upregulation of Il6, Il1b, Tnf, and Ifng, as well as the downregulation of Pparg and Il10 (Fig. 2d). Furthermore, to reveal common or distinct regulation of gene annotation categories in the transcriptome or proteome we performed a two dimensional annotation enrichment analysis^23^ (Table S4). Again, most gene categories regulated by age were showing the same direction in transcriptome and proteome so that the positive Pearson correlation of the annotation enrichment scores was highly significant (Fig. 2e). We observed several hallmarks of aging, including a decline in mitochondrial function and upregulation of pro-inflammatory pathways (‵inflammaging′). Interestingly, we detected a strong increase in immunoglobulins in both datasets, as well as higher levels of MHC class I, which is consistent with the observed increase in the interferon pathway (Fig. 2e). Many extracellular matrix genes, such as Collagen III were downregulated on both the mRNA and protein level (Fig. 2f), while the levels of all basement membrane-associated Collagen IV genes were increased on the protein level, but decreased at the mRNA level (Fig. 2g). The differential regulation of Collagen IV transcripts and proteins highlights the importance of combined RNA and protein analysis. We validated the increased protein abundance of Collagen IV using immunofluorescence and found that interestingly the main increase in Collagen IV in old mice was found around airways and vessels (Fig. 2h).

The combination of tissue proteomics with single cell transcriptomics enabled us to predict the cellular source of the regulated proteins, which can be explored in the online webtool. Furthermore, single cell RNAseq can disentangle relative frequency changes of cell types from real changes in gene expression within a given cell type. We analyzed age dependent alterations of relative frequencies of the 30 cell types represented in our dataset. Since the cell type frequencies are proportions the data is compositional. Therefore, it is impossible to statistically discern if a relative change in cell type frequency is caused by the increase of a given cell type or the decrease of another. However, after performing dimension reduction using multidimensional scaling of the cell type proportions we observed a significant association between the first coordinate and age (Fig. S3a, b; Wilcoxon test, P < 0.005), indicating that cell type frequencies differed between young and old mice. Interestingly, the ratio of club to ciliated cells was altered in old mice. While Dropseq data from young mice contained more club cells than ciliated cells, this ratio was inverted in old mice with more than twofold more ciliated cells over Club cells (Fig. S3c). We validated this finding *in situ* by quantifying airway club and ciliated cells using immunostainings of Foxj1 (ciliated cell marker) and CC10 (club cell marker) (Fig. S3d). Club cell numbers were decreased in old mice (Fig S3e), while ciliated cells were increased (Fig. S3f), leading to a significantly altered ratio of club to ciliated cells in aged mouse airways (Fig. S3g).

### Age dependent changes in composition and organization of the pulmonary extracellular matrix

The extracellular matrix (ECM) can act as a solid phase-binding interface for hundreds of secreted proteins, creating an information-rich signaling template for cell function and differentiation^24^. Alterations in ECM composition and possibly architecture in the aging lung have been suggested^25^, however experimental evidence using unbiased mass spectrometry is scarce. We previously developed the quantitative detergent solubility profiling (QDSP) method to add an additional dimension of protein solubility to tissue proteomes^26–28^. In QDSP, proteins are extracted from tissue homogenates with increasing stringency of detergents, which typically leaves ECM proteins enriched in the insoluble last fraction. This enables better coverage of ECM proteins and analysis of the strength of their associations with higher order ECM structures such as microfibrils or collagen networks. We applied this method to young and old mice and compared protein solubility profiles between the two groups (Fig. 3a). Differential comparison of the solubility profiles between young and old mice revealed 74 proteins, including eight ECM proteins, with altered solubility profiles (FDR <20%) (Table S5).

**Figure 3.**
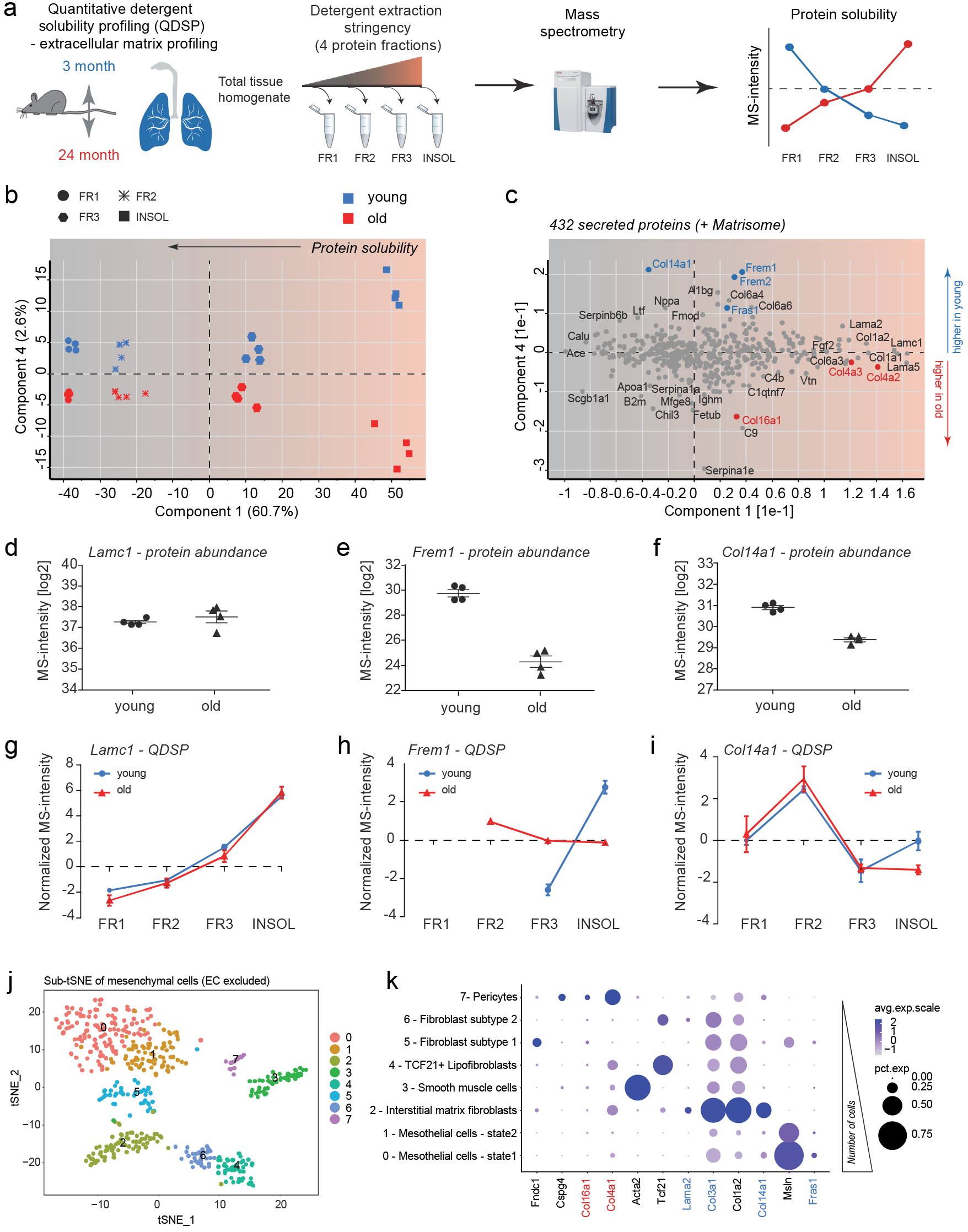
Quantitative detergent solubility profiling of the aging lung proteome. (a) Experimental design – extraction of proteins from whole lung homogenates with increasing detergent stringency results in four distinct protein fractions, which are analyzed by mass spectrometry. (b) The projections of a principal component analysis (PCA) of 432 proteins with the annotation ‵secreted′ in the Uniprot and/or Matrisome database separate the four protein fractions, indicated by symbol shape, in component 1 and the age groups, as indicated by color, in component 4. (c) The loadings of the PCA are shown. (d-f) Relative differences in MS-intensity (abundance) of the indicated proteins. (g-i) The normalized MS-intensity across the four protein fractions from differential detergent extraction highlights changes in protein solubility between young and old mice for the indicated proteins. (j) The tSNE map shows 483 cells of mesenchymal origin (endothelial cells excluded), revealing 8 distinct cellular states. (k) The dotplot shows (1) the percentage of cells expressing the respective selected marker gene using dot size and (2) the average expression level of that gene based on UMI counts. Proteins that were found to be up-or downregulated are highlighted in red and blue respectively.

Using principal component analysis of 432 secreted extracellular proteins we found that the protein solubility fractions separated in component 1, while the age groups separated in component 4 of the data (Fig. 3b). Thus, principal component analysis enabled the stratification of secreted proteins by their biochemical solubility and their differential behavior upon aging (Fig. 4c). This analysis also showed that neither the abundance nor the solubility of many ECM proteins, including Collagen I and basement membrane laminins, was altered (Fig. 3c). We uncovered several ECM proteins with greatly reduced abundance and/or significant changes in their solubility profiles. While the most abundant basement membrane laminin chain (Lamc1) was unaltered in both abundance (Fig. 3d) and solubility (Fig. 3g), serving as a control for overall integrity of the basement membrane and the quality of our data, the basement membrane-associated trimeric Fraser Syndrome complex (consisting of Fras1, Frem1, and Frem2) was downregulated (Fig. 3e) and more soluble (Fig. 3h) in old age.

**Figure 4.**
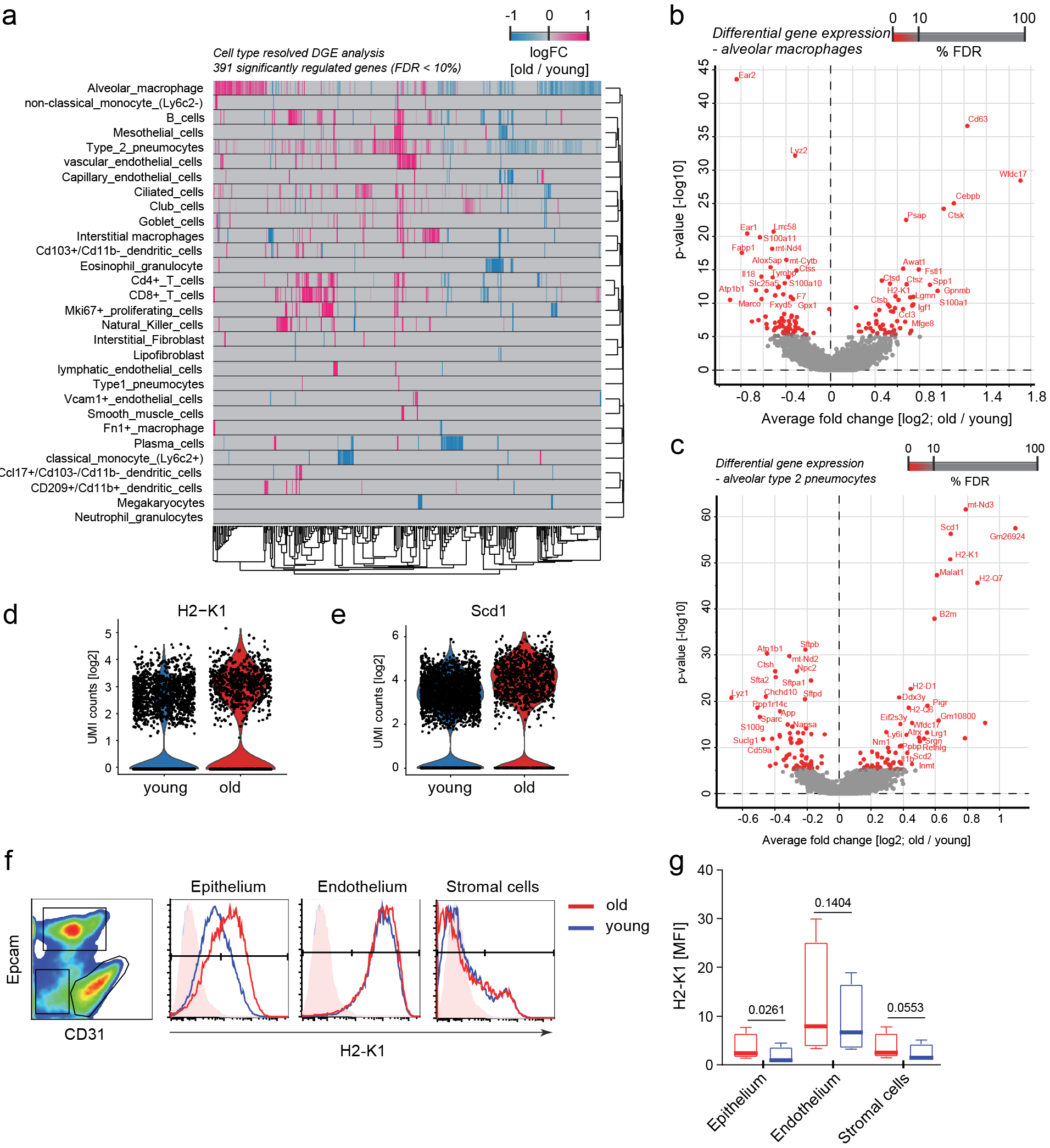
Cell type specific differential gene expression analysis. (a) Heatmap displays fold changes derived from the cell type resolved differential expression analysis. Rows and columns correspond to cell types and genes, respectively. Negative fold change values (blue) represent higher expression in young compared to old. Positive fold change values are colored in pink. (b, c) Volcano plots visualize the differential gene expression results in Alveolar macrophages (b) and alveolar type-2 pneumocytes (c). X and Y axes show average log2 fold change and −log10 p-value, respectively. (d, e) Violin plots depict normalized expression values of two exemplary genes H2-K1 (d) and Scd1 (e) across age groups for type-2 pneumocytes. (f) The indicated cell lineages were gated by flow cytometry as shown in the left panel in a CD31 and Epcam co-staining and evaluated for H2-K1 expression on protein level. The histograms show fluorescence intensity distribution of the H2-K1 cell surface staining for the indicated lineages and age groups. (g) Boxplot shows mean fluorescence intensity for H2-K1 in the indicated cell lineages across four replicates of mice. P-values are from an unpaired, two-tailed t-test.

Incorporation of the Fraser syndrome complex within the basement membrane (rendering it more insoluble) has been shown to depend on extracellular assembly of all three proteins^29^, indicating that this assembly and/or the expression of either one or all subunits of the complex is perturbed in old mice. Fraser syndrome is a skin-blistering disease which points to an important function of the Fraser syndrome complex proteins in linking the epithelial basement membrane to the underlying mesenchyme^29^. In the lungs of adult mice, expression is restricted to the mesothelium; Fras1 −/− mice develop lung lobulation defects^30^. Indeed, we were able to confirm the mesothelium specific expression of Fras1 using our scRNAseq data (Fig. 3j, k). Another ECM protein with markedly reduced protein levels (Fig. 3f) and increased solubility (Fig. 3l) was Collagen XIV, a collagen of the FACIT family of collagens that is associated with the surface of Collagen I fibrils and may function by integrating collagen bundles^31^. Collagen XIV is a major ECM binding site for the proteoglycan Decorin^32^, which is known to regulate TGF-beta activity^33, 34^. Interestingly, our scRNAseq data localized Collagen XIV expression to interstitial fibroblasts, which also specifically expressed Decorin and were distinct from the lipofibroblasts that showed very little expression of this particular collagen (Fig. 3 j/k).

### Cell type specific effects of aging

Cell type-resolved differential gene expression testing between age groups in the single cell data sets identified 391 significantly regulated genes (FDR < 10%) (Fig. 4a; Table S6). Alveolar macrophages and type 2 pneumocytes, the two cell types with highest number of cells in the dataset, are discussed as an example for the type of insight that can be gained from our cell type resolved resource. Both cell types showed a clearly altered phenotype in aged mice.

In alveolar macrophages, we found 125 significantly regulated mRNAs (FDR<10%, Fig. 4b), including the downregulation of the genes for Eosinophil cationic protein 1 & 2 (Ear1 & Ear2), which have ribonuclease activity and are thought to have potent innate immune functions as antiviral factors^35^. The cell surface scavenger-type receptor Marco, mediating ingestion of unopsonized environmental particles^36^, was downregulated in old mice which may cause impaired particle clearance by alveolar macrophages. We observed higher levels of the C/EBP beta (Cebpb), which is an important transcription factor regulating the expression of genes involved in immune and inflammatory responses^37, 38^. Several genes that have been shown to be upregulated in lung injury, repair and fibrosis^28^, such as Spp1, Gpnmb, and Mfge8 were also induced in alveolar macrophages of old mice, which may be a consequence of the ongoing ‵inflammaging′.

In alveolar type 2 pneumocytes, 121 mRNAs were significantly regulated (FDR<10%, Fig. 4c). We observed a strong increase of the Major Histocompatibility Complex (MHC) class I proteins H2-K1, H2-Q7, H2-D1, and B2m (Fig. 4c, d), which we validated using an independent flow cytometry experiment (Fig. 4f). Elevated MHC class I levels likely result in increased presentation of self-antigens to the immune system and are consistent with our observation of a prominent Interferon-gamma signature in old mice (Fig. 2d), which is known to activate MHC class I expression^39^. Type-2 pneumocytes of old mice featured a highly significant upregulation of the enzyme Acyl-CoA desaturase 1 (Scd1), which is the fatty acyl Δ9-desaturating enzyme that converts saturated fatty acids (SFA) into monounsaturated fatty acids (MUFA) (Fig. 4c, e). The age dependent upregulation of Scd1 in type-2 pneumocytes is a novel observation, which may have important implications since Scd1 is thought to induce adaptive stress signaling that maintains cellular persistence and fosters survival and cellular functionality under distinct pathological conditions^40^.

To obtain a meta-analysis of changes in previously characterized gene expression modules and pathways, we used cell type resolved mRNA fold changes for gene annotation enrichment analysis (Fig. 5a and b, Table S7) and upstream regulator analysis (Fig. 5c-e). The analysis revealed cell type specific alterations in gene expression programs upon aging. For instance, comparing club cells to type-2 pneumocytes, showed that Nrf2 (Nfe2l2) mediated oxidative stress responses were higher in type-2 pneumocytes of old mice and lower in club cells (Fig. 5c). Aging is known to affect growth signaling via the evolutionary conserved Igf-1/Akt/mTOR axis^2^. Interestingly, we found evidence for increased mTOR signaling in type-2 and club cells, but not in ciliated and goblet cells (Fig. 5c). Mesenchymal cells showed remarkable differences in their aging response (Fig. 5d). For instance, we observed the pro-inflammatory Il1b signature in capillary endothelial cells, as well as in mesothelial and smooth muscle cells, but not in the other mesenchymal cell types. In myeloid cell types we found both differences and similarities in the aging response (Fig. 5e). While, an increased interferon gamma and reduced Il10 signature in old mice was consistently observed, other effects were more specific, such as the increase in Stat1 target genes in classical monocytes (Ly6c2+), which was not observed in non-classical monocytes(Ly6c2-).

**Figure 5.**
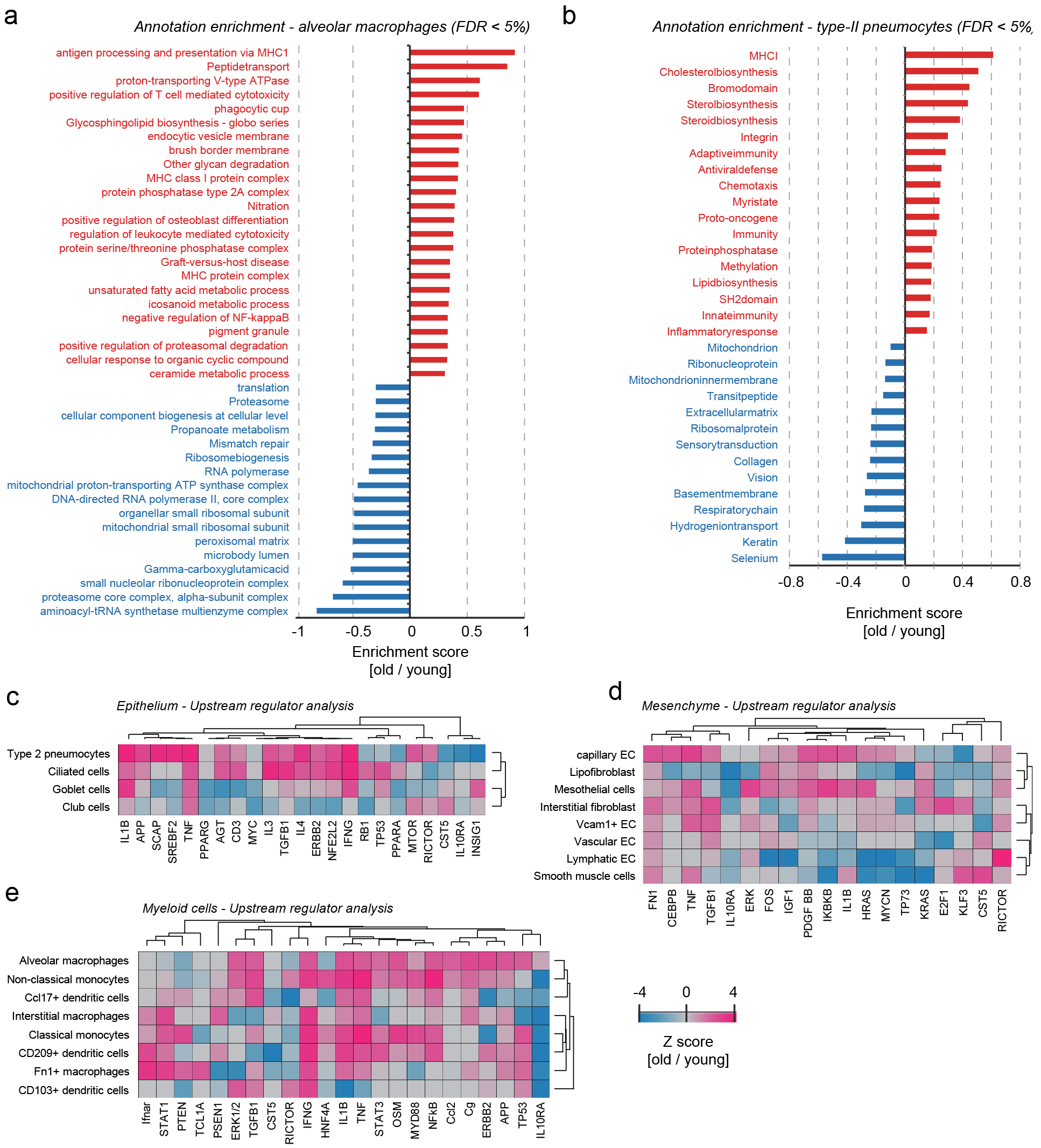
Cell type specific pathway and upstream regulator analysis. (a, b) The bar graph shows the result of a gene annotation enrichment analysis for (a) alveolar macrophages, and (b) type-II pneumocytes, respectively. Gene categories with positive (upregulated in old) and negative scores (downregulated in old) are highlighted in red and blue respectively. (c-e) Upstream regulators are predicted based on the observed gene expression changes for (c) epithelial, (d) mesenchymal, and (e) myeloid cells. Cell types and regulators were grouped by unsupervised hierarchical clustering (Pearson correlation) and the indicated transcriptional regulators and cytokines, growth factors and ECM proteins are color coded based on the activation score as shown.

### Increased cholesterol biosynthesis in aged type-2 pneumocytes and lipofibroblasts

Pulmonary surfactant homeostasis is a tightly regulated process that involves synthesis of lipids by type-2 pneumocytes and lipofibroblasts^41^. Lipid metabolism in alveolar type-2 cells is regulated by sterol-response element-binding proteins (SREBPs), such as Srebf2 and their negative regulators Insig1 and Insig2. Deletion of Insig1/2 in mouse type-2 pneumocytes activated SREBPs and led to the accumulation of neutral lipids (cholesterol esters and trigylcerids) in type-2 pneumocytes and alveolar macrophages, accompanied by lipotoxicity-related lung inflammation and tissue remodeling^42^. Interestingly, we observed very similar gene expression changes in type-2 pneumocytes of old mice as reported for the Insig1/2 deletion. Consistently, the upstream regulator analysis predicted increased activity of Srebf2 and reduced activity of Insig1 specifically in type-2 pneumocytes of old mice (Fig. 5c). The upstream regulator analysis was based on 25 known targets of SREBP/Insig1, all of which were increased in aged type-2 pneumocytes (Fig. 6a). Using gene annotation enrichment analysis on Uniprot Keywords, GO terms and KEGG pathways (Table S7) we found increased cholesterol biosynthesis as the top hit in type-II pneumocytes and lipofibroblasts and no other cell type (Fig. 6b). Indeed, most of the Insig1/2 target genes are directly involved in cholesterol biosynthesis (Fig. 6c).

**Figure 6.**
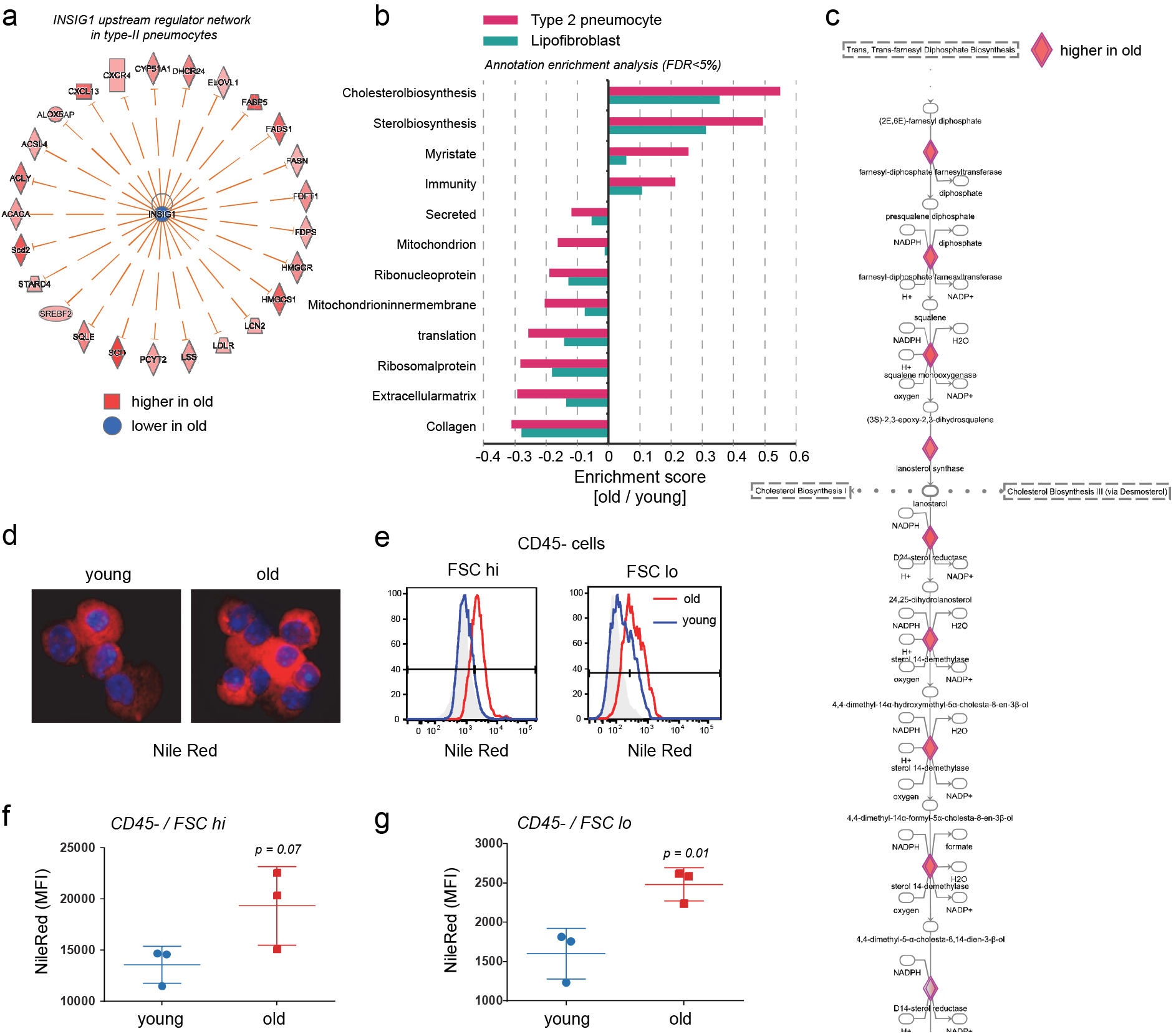
Increased cholesterol biosynthesis in type 2 pneumocytes and lipofibroblasts in aged mice. (a) The graph shows genes known to be negatively regulated by Insig1 that were found to be upregulated in type-2 pneumopcytes of old mice. (b) Selected gene categories found to be significantly (FDR <5%) up-(positive enrichment scores) or downregulated (negative enrichment scores) in the indicated cell-types. (c) Segment of the cholesterol biosynthesis pathway. Diamant shaped nodes represent enzymes that were found to be upregulated in type-2 pneumocytes of old mice. The biochemical intermediates are named in between the enzyme nodes. (d) Nile red staining of neutral lipids in cells from a whole suspension depleted for CD45+ leukocytes (CD45 lineage negative cells). (e) Quantification of nile red stainings using flow cytometry. Histograms show flow cytometry analysis of Nile Red in aged (red) and young (blue) mice, unstained control is represented in gray. Cells were stratified by size in bins of large (FSC hi) and small (FSC lo) cells using the forward scatter. (f, g) Nile Red mean fluorescence intensity (MFI) quantification across three individual mice for (f) CD45 negative and forward scatter (FSC) high, and (g) FSC low cells. P-values are from an unpaired, two-tailed t-test.

To confirm the increased cholesterol biosynthesis and analyze the actual lipid content of the cells we used the Nile red dye to stain neutral lipids in cells of a whole lung suspension after depletion of leukocytes (Fig. 6d). Using flow cytometry we quantified the Nile red lipid staining and found a significant increase in both mean fluorescence intensity (Fig. 6d, e), and number of Nile red positive cells (Fig. 6f, g) in the CD45 negative cells of old mice. CD45+ cells were not significantly altered, indicating that the increase in neutral lipid content is specific to epithelial cells and fibroblasts. Thus, we have shown for the first time that increased cholesterol biosynthesis and neutral lipid content in type-2 pneumocytes and lipofibroblasts is a hallmark of lung aging.

## Discussion

Increasing both health and lifespan of humans is one of the prime goals of the modern society. In order to better understand age-related chronic lung diseases such as COPD, lung cancer or fibrosis, intense efforts in integrated multi-omics systems biology tools for the analysis of lung aging are needed^25^. In this work, we present the first single cell survey of mouse lung aging and computationally integrate transcriptomics data with deep tissue proteomics data to build an atlas of the aging lung. The lung aging atlas and associated raw data can be accessed at https://theislab.github.io/LungAgingAtlas (Figure S4). It features five dimensions that can be navigated through gene and cell type specific queries: (1) cell type specific expression of genes and marker signatures for 30 cell types, (2) regulation of gene expression by age on cell type level, (3) cell type resolved pathway and gene category enrichment analysis, (4) regulation of protein abundance by age on tissue level, (5) regulation of protein solubility by age.

The highly multiplexed nature of droplet based single cell RNA sequencing used in this study allows the direct analysis of thousands of individual cells freshly isolated from whole mouse lungs, providing unbiased classification of cell types and cellular states. Two previous studies have analyzed aging effects using single cell transcriptomics^20, 21^ and found increased transcriptional variability between cells in human pancreas and T cells. In this study, we identify aging associated increased transcriptional noise, which may result from deregulated epigenetic control, in most cell types of the lung, indicating that this phenomenon is a general hallmark of aging that likely affects most cell types in both mice and humans. This new concept is supported by our study and it will be interesting when and how future investigations will shed light on the molecular mechanisms driving this phenomenon.

We have used two independent cohorts of young and old mice and uncovered remarkably well conserved aging signatures in both mRNA and protein. Thus, the two datasets validate each other and show that on gene category and pathway level, the analysis of protein and mRNA content can lead to overall similar results with important differences. Hallmarks of aging, such as the downregulation of mitochondrial oxidative phosphorylation and the upregulation of pro-inflammatory signaling pathways were consistently observed in both datasets. On the level of individual genes/proteins, however, we often observed interesting differences, which indicates that for functional analysis of a particular gene/protein it remains essential to also analyze the protein, which ultimately executes biological functions.

The example of basement membrane collagen IV genes that were all downregulated on the mRNA level but upregulated on the protein level serves as a reminder that mRNA analysis should always be taken with a grain of salt. Next to mass spectrometry based methods, novel single cell methods combining mRNA and protein analysis, such as CITE-seq^43^, will become ever more important in the near future. We show that the combination of single cell-resolved mRNA analysis and bulk proteomics is highly complementary by using the single cell expression data to understand the most likely cellular origin of proteins that showed altered abundance with age. Spatial transcriptomics methods for high throughput detection of transcripts in single cells *in situ* are currently quickly evolving^44, 45^. Traditional antibody based methods for single cell protein analysis *in situ* are however not well multiplexed and do not easily scale for high throughput. Thus, to fully develop the enormous potential of single cell multi-omics data integration (M.Colomé-Tatché … F.J.Theis 2018; https://doi.org/10.1016/j.coisb.2018.01.003), the field depends on current and future developments in methods for simultaneous protein and mRNA analysis on single cell level *in situ*^46, 47^.

We analyzed the foundations of lung tissue architecture by quantifying compositional and structural changes in the aged extracellular matrix using state of the art mass spectrometry workflows. The ECM is not only key as a scaffold for the lungs overall architecture, but also an important instructive niche for cell fate and phenotype^24, 48^. Recent proteomic studies have demonstrated that at least 150 different ECM proteins, glycosaminoglycans and modifying enzymes are expressed in the lung, and these assemble into intricate composite biomaterials that are characterized by specific biophysical and biochemical properties^27, 28, 49^. Due to this complexity of the ECM, both in terms of composition and posttranslational modification, and the assembly of ECM proteins into supramolecular structures, it is presently unclear on which level and how exactly the aging process affects the lungs′ ECM scaffold. We used detergent solubility profiling to screen for differences in protein crosslinking and complex formation within the ECM. Surprisingly, most solubility profiles were not significantly altered with age, indicating that aging-related ECM remodeling does not involve large differences in covalent protein crosslinks. However, we observed a few very strong changes in the ECM, which were completely novel in the context of aging and are open for future investigation into their functional implications.

In order to stabilize the alveolar structure during breathing-induced expansion and contraction, type-2 pneumocytes produce and secrete pulmonary surfactant, which is a thin film of phospholipids and surfactant proteins^41^. The lipid composition of pulmonary surfactant has been shown to change with age^50^, and electron microscopy of surfactant and the lipid loaded lamellar bodies in type-2 pneumocytes revealed ultrastructural disorganization with age^51^. This may be related to our finding that cholesterol biosynthesis and neutral lipid content is upregulated in type-2 cells of old mice. It is currently unclear at which level the homeostasis of lipid metabolism is altered in the aged lungs. We found strong similarity of the aged type-2 phenotype with the phenotype in Insig1/2 knock out mice that accumulated neutral lipids, accompanied by lipotoxicity-related lung inflammation and tissue remodeling^42^. Thus, it is possible that part of the chronic inflammation we observed in the aged lung is influenced by deregulation of lipid homeostasis. The inflammatory phenotype may also be related to epithelial senescence, as mice with a type-2 pneumocyte specific deletion of telomerase, and thus premature aging with increased senescence in these cells, developed a pro-inflammatory tissue microenvironment and were less efficient in resolving acute lung injury^52^.

In summary, we have demonstrated that the lung aging atlas presented here contains a plethora of novel information on molecular and cellular scale and serves as a reference for the large community of scientists studying chronic lung diseases and the aging process. The data also serves as analysis template for the currently ongoing worldwide collection of data for a Human Lung Cell Atlas, which will hopefully enable analysis of aging effects in human lungs.

## Supplementary data and Methods

**Supplementary Figure S1.**
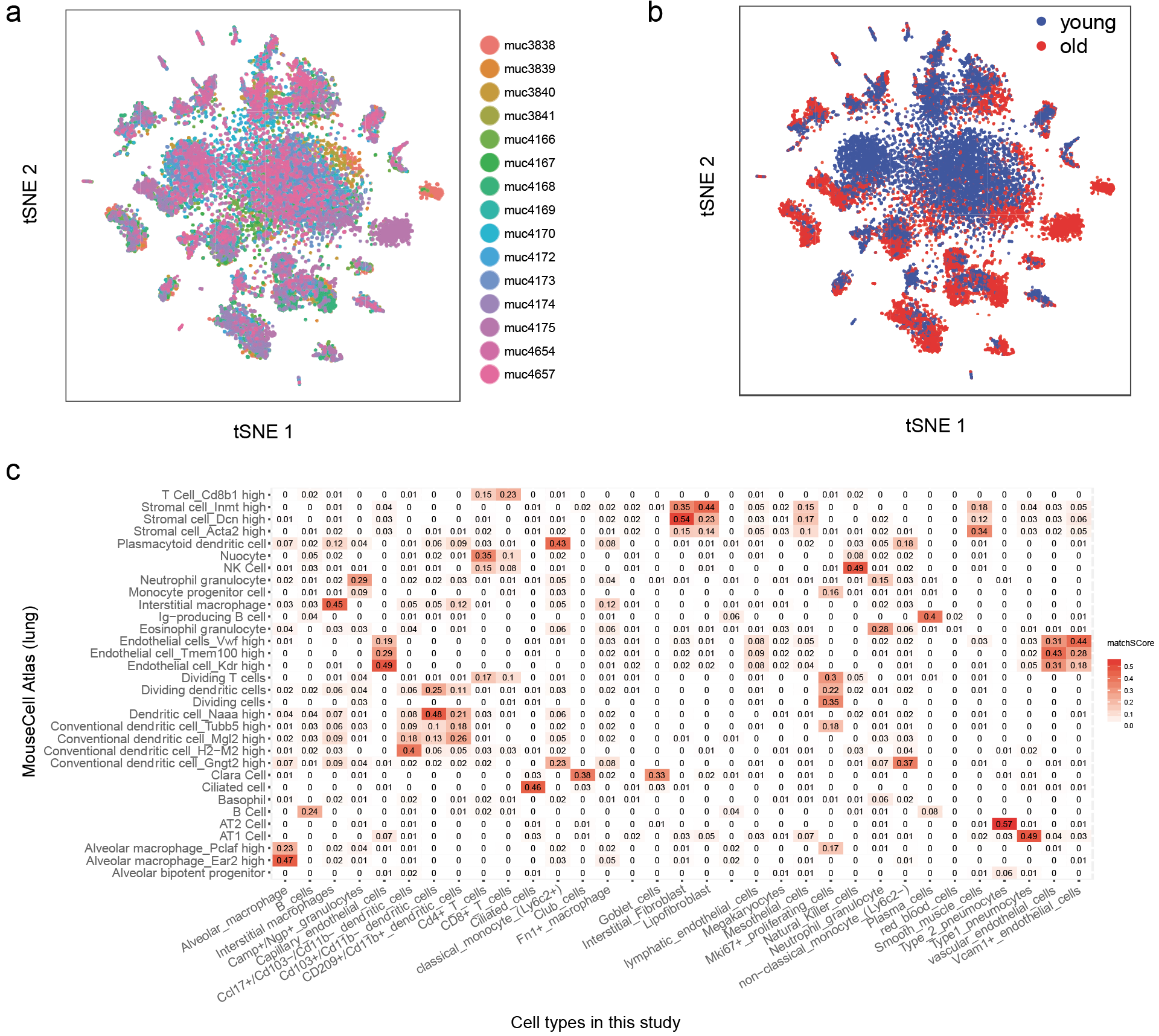
Excellent technical reproducibility and correct cell type annotation. (a, b) tSNE visualization colored by mouse sample (a) and age group (b). machSCore visualization of overlap between cell type markers derived from the Mouse Cell Atlas (lung) and this study (c). Red colors indicate higher matchSCore values.

**Supplementary Figure S2.**
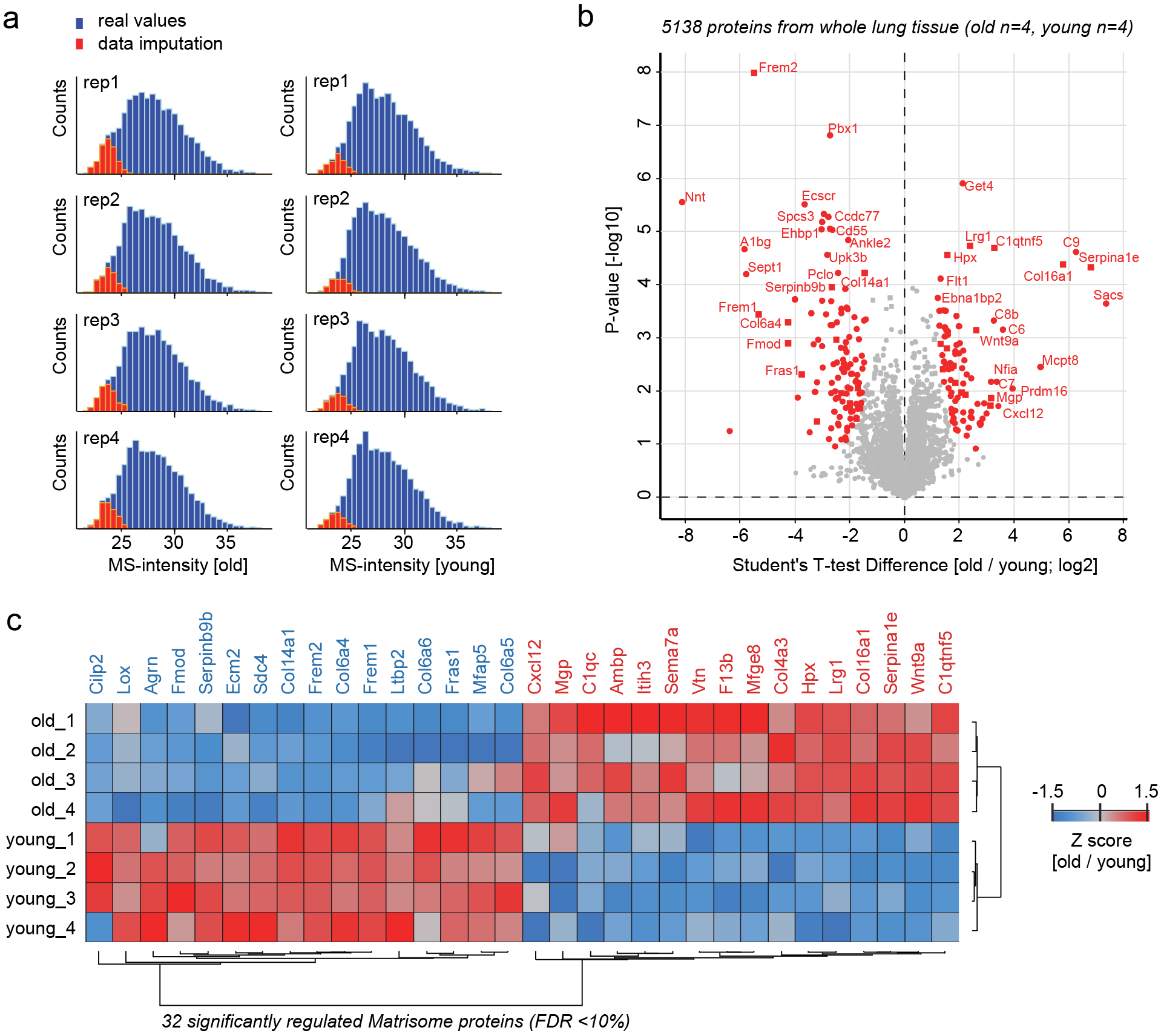
Whole lung tissue proteomes of mouse lung aging. (a) The histogram shows the normal distribution of mass spectrometry (MS) intensities for the quantified proteins in all replicates and experimental groups. For t-test statistics the missing values (proteins not detected in the respective sample) were replaced by a normal distribution of random values at the lower end of the distribution of real values (data imputation). (b) Proteins regulated with a false discovery rate < 10% are highlighted in red in the vulcano plot showing the indicated fold changes and p-values from t-test statistic. Matrisome proteins are shown as rectangles and all other proteins as filled circles. (c) The z-score values of 32 significantly regulated extracellular matrix proteins were grouped by unsupervised hierarchical clustering (Pearson correlation)

**Supplementary Figure S3.**
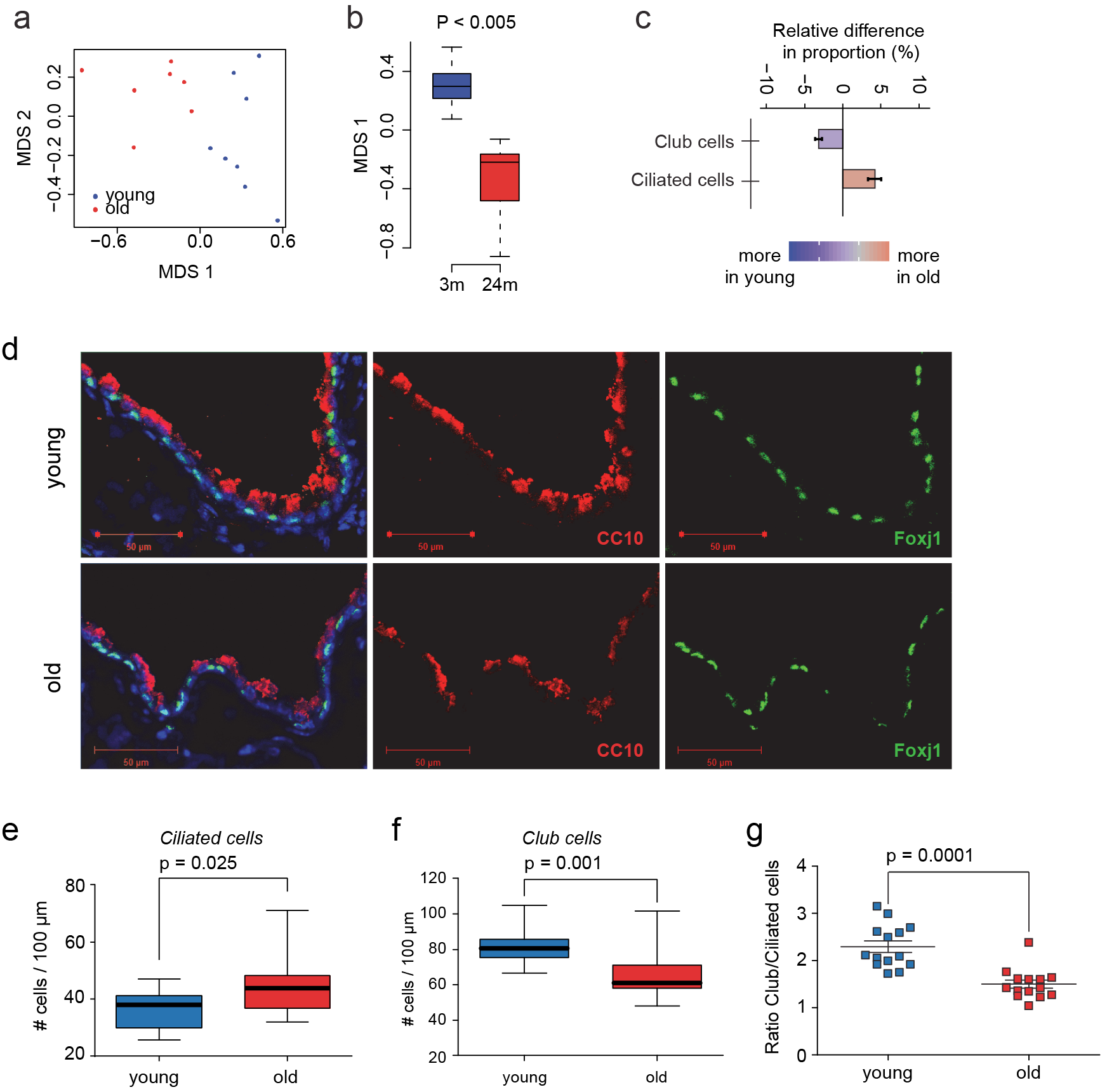
Cell type frequency analysis reveals an altered ratio of club to ciliated cells in airways of old mice. (a) The MDS plot shows the mouse-wise euclidean distances of cell type proportions for the two age groups (b) The box plot shows the significant difference in euclidean distance of cell type proportions. (c) Relative difference in proportion in percent of all cell in the dataset for the indicated cell types. (d) Club and ciliated cells were stained using a CC10 and Foxj1 antibody respectively. (e, f) The box plots show the quantification of (e) Ciliated and (f) Club cells from counting 2647 cells in 14 airways of two mice of each age group. (g) Ratio of ciliated to club cells in 14 airways. P-values are from an unpaired, two-tailed t-test using Welch′s correction.

**Supplementary Figure S4.**
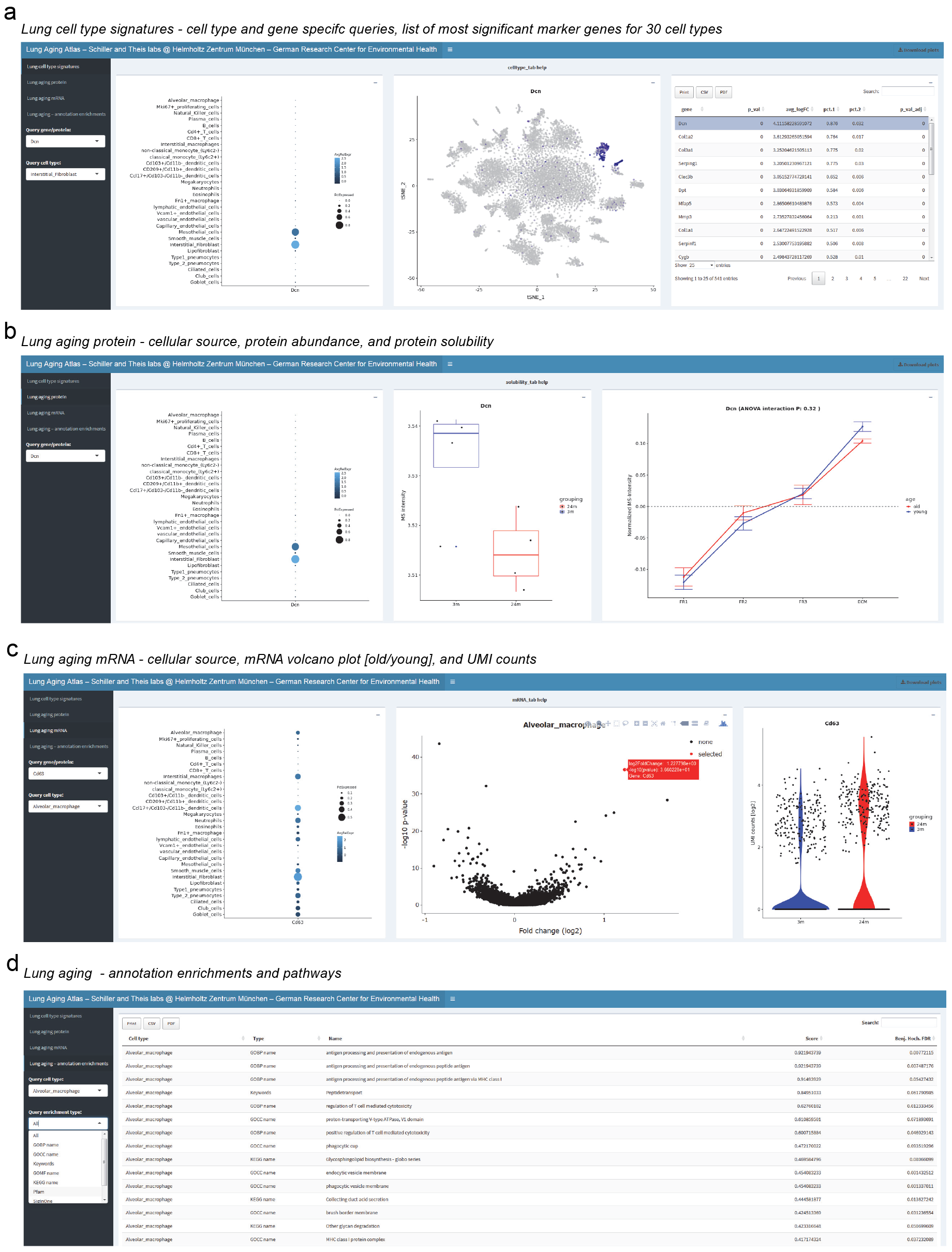
A user friendly and interactive webtool to navigate the Lung Aging Atlas. (a) The first tab ‵Lung cell type signatures′ provides a cell type dotplot (left panel) and a color coded tSNE map (middle panel) for gene specific queries and illustrates cell type specific expression of any gene of interest. A cell type query produces a list of top marker genes for the cell type of interest (right panel). (b) The panel ‵Lung aging protein′ features a dot plot to illustrate the most likely cellular source of the protein of interest (left panel), a box plot to show alterations in total lung tissue protein abundance in old mice (middle panel), and a line plot to show protein solubilities. Protein solubility is measured by relative quantification of protein abundance across four fractions. Fraction 1 (FR1) contains proteins with highest and fraction 4 (ECM) with lowest solubility. Curves that peak on the right (ECM) thus represent insoluble proteins. (c) The tab ‵Lung aging mRNA′ again features the dotplot (left panel), a volcano plot that shows fold changes [old/young] on the x-axis and −log10 p-values on the y-axis (middle panel), and a violin plot of the log2 UMI counts illustrating mRNA abundance in young and old mice. The dot and violin plot are navigated with the gene specific query, while the volcano plot requires navigation via the cell type query. The volcano plot has a toggle over function that allows identification of genes and can thus be used to browse through differential gene expression between young and old cells of any cell type of interest. (d) In the tab ‵Lung aging-annotation enrichments′, the gene annotation enrichments between old and young can be browsed for all 30 cell types.

## Methods

### Generation of single cell suspensions from whole mouse lung

Lung tissue was perfused with sterile saline from the right to the left ventricle of the heart and subsequently inflated via a catheter in the trachea, by an enzyme mix containing dispase (50 caseinolytic units/ml), collagenase (2 mg/ml), elastase (1 mg/ml), and DNase (30 ug/ml). After tying off the trachea, the lung was removed and immediately minced to small pieces (approx. 1 mm^2^). The tissue was transferred into 4 ml of enzyme mix for enzymatic digestion for 30 min at 37°C. Enzyme activity was inhibited by adding 5 ml of PBS supplemented with 10% fetal calf serum. Dissociated cells in suspension were passed through a 70 μm strainer and centrifuged at 500 × g for 5 min at 4°C. Red blood cell lysis (Thermo Fischer 00-4333-57) was done for 2 min and stopped with 10% FCS in PBS. After another centrifugation 5 min at 500 × g (4°C) the cells were counted using a Neubauer chamber and critically assessed for single cell separation and viability. 250.000 cells were aliquoted in 2.5 mL of PBS supplemented with 0.04% of bovine serum albumin and loaded for Drop-Seq at a final concentration of 100 cells/μL.

### Single cell RNA sequencing

Drop-seq experiments were performed largely as described previously^15, 16, 53^, with few adaptations during the single cell library preparation. Briefly, using a microfluidic PDMS device (Nanoshift), single cells (100/μl) from the lung cell suspension were co-encapsulated in droplets with barcoded beads (120/μl, purchased from ChemGenes Corporation, Wilmington, MA) at rates of 4000 μl/hr. Droplet emulsions were collected for 15 min/each prior to droplet breakage by perfluorooctanol (Sigma-Aldrich). After breakage, beads were harvested and the hybridized mRNA transcripts reverse transcribed (Maxima RT, Thermo Fisher). Unused primers were removed by the addition of exonuclease I (New England Biolabs), following which beads were washed, counted, and aliquoted for pre-amplification (2000 beads/reaction, equals ~100 cells/reaction) with 12 PCR cycles (primers, chemistry, and cycle conditions identical to those previously described^15^. PCR products were pooled and purified twice by 0.6× clean-up beads (CleanNA). Prior to tagmentation, cDNA samples were loaded on a DNA High Sensitivity Chip on the 2100 Bioanalyzer (Agilent) to ensure transcript integrity, purity, and amount. For each sample, 1 ng of pre-amplified cDNA from an estimated 1000 cells was tagmented by Nextera XT (Illumina) with a custom P5 primer (Integrated DNA Technologies). Single cell libraries were sequenced in a 100 bp paired-end run on the Illumina HiSeq4000 using 0.2 nM denatured sample and 5% PhiX spike-in. For priming of read 1, 0.5 μM Read1CustSeqB (primer sequence: GCCTGTCCGCGGAAGCAGTGGTATCAACGCAGAGTAC) was used.

### Bioinformatic processing of scRNA-seq reads

The Dropseq core computational pipeline was used for processing next generation sequencing reads of the scRNA-seq data, as previously described^15^. Briefly, STAR (version 2.5.2a) was used for mapping^54^. Reads were aligned to the mm10 genome reference (provided by Drop-seq group, GSE63269). For barcode filtering, we excluded barcodes with less than 200 genes detected.

### Single cell data analysis

After constructing the single cell gene expression count matrix, we used the R package *Seurat^55^* and custom scripts for analysis.

#### Quality control

A high proportion of transcript counts derived from mitochondria-encoded genes can indicate low cell quality. Similarly cells with unusually high UMI count levels may represent cell doublets. Therefore we removed cells with >10% mitochondria derived counts and >5000 total UMI counts from the downstream analysis.

#### Unsupervised clustering and visualization

Highly variable genes were defined within each mouse sample (young, n=8; old n=7) separately following Seurat standard approach. Next, genes appearing in >4 mouse samples in the set of highly variable genes were defined as a set of consensus highly variable genes. To minimize the effect of cell-cycle on clustering we removed cell-cycle genes^56^ from the set of consensus highly variable genes. All 14813 cells passing quality control were merged into one count matrix and normalized and scaled using Seurat’s NormalizeData() and ScaleData() functions. The reduced set of consensus highly variable genes was used as the feature set for independent component analysis using Seurat’s RunICA() function. The first 50 independent components were used for tSNE visualization and Louvain clustering using the Seurat functions RunTSNE() and FindClusters(), respectively.

#### Cell type marker discovery

The Seurat FindAllMarkers() function was used to identify cluster specific marker genes. Based on manual annotation and with guidance of the enrichment analysis (see below) the 36 clusters were assigned to 30 cell type identities. Using the annotation of cell type identities the FindAllMarkers() function was called to identify the final set of cell type markers used throughout this analysis.

#### Ambient RNA identification

An important technical detail needed our attention and is briefly described here. As infrequently discussed in the community but not yet addressed, we also observed ‵ambient mRNA′ effects, which we believe are the consequence of free mRNA released from dying cells hybridizing with beads in droplets during the microfluidic capture of single cells in the Dropseq workflow. The ambient mRNAs are typically derived from highly abundant transcripts and this artefact is inherent to all droplet based methods (including the commercially available 10x platform). Here, it can be exemplified by the Scgb1a1 gene in Figure 1C that is known to be highly specific for Club and Goblet cells but was nevertheless detected in almost 100% of the cells in our data. However, the UMI count levels were much higher in Club and Goblet cells (representing the real source of expression) indicating that the mRNA counts observed in all other clusters were ambient mRNA background. To independently confirm this we therefore determined all genes that showed ambient mRNA background by analyzing the identity of genes on beads at the tailend of the total UMI count distribution (on average 10 UMIs per barcode), representing empty beads that were never in contact with a real cell but nevertheless contain information from free floating ambient mRNA. We identified 153 genes (Table S8) with an ‘ambient mRNA’ effect and accounted for this effect in downstream analysis.

#### Cell type annotation

To aid the assignment of cell type to clusters derived from unsupervised clustering, we performed cell type enrichment analysis. Cell type gene signatures obtained from bulk level gene expression were downloaded from the ImmGen and xCell resources. Each gene signature obtained from our clustering was statistically evaluated for overlap with gene signatures contained in these two resources. Mouse gene symbols were capitalized to map to human gene symbols. Overlap between gene signatures was evaluated using Fisher’s Exact test.

#### Mouse Cell Atlas integration

Cell type marker signatures in our data (Table S1) were compared to cell type marker signatures in the Mouse Cell Atlas (MCA)^13^. MatchSCore (Mereu et al 2018, bioRxiv doi: https://doi.org/10.1101/314831) was used to quantify overlap between cell type marker signatures derived from our study and the MCA. Marker genes with adjusted p-value < 0.1 and average log fold change > 1 were considered.

#### Quantifying transcriptional noise

Transcriptional noise in the gene expression profiles was quantified in a multivariate fashion following previous work^21^. Briefly, the euclidean distance between the gene expression profile of each cell and the average expression profile of the respective cell type was calculated for each mouse. Median euclidean distance was used to summarize transcriptional noise at the mouse level. To statistically assess the association between transcriptional noise and age within each cell type Wilcoxon’s rank sum test was used. To statistically assess the global association between transcriptional noise and age, analysis of variance was used In particular, transcriptional noise was modelled as the dependent variable with cell type as a covariate and age as the explanatory variable.

#### Cell type frequency analysis

Cell type frequencies were calculated based on the counts of cells annotated to each cell type for each mouse. Counts were transformed to proportions using the DR_data() function of the DirichletReg R package which causes the values to shrink away from extreme values of 0 and 1. Next, the mouse-wise euclidean distances were calculated based on these proportions using the dist() R function followed by multidimensional scaling using the isoMDS() R function. To statistically assess the association between age and the first coordinate derived from the multidimensional scaling Wilcoxon test was applied. Relative changes in cell type frequencies were calculated by subtracting the median cell type proportion of the young mice from the cell type proportions of the old mice.

#### Cell type resolved differential expression analysis

Cell type resolved differential expression analysis was performed using the Seurat differential gene expression testing framework. Within each cell type cells were grouped by age and differential testing performed using the Seurat FindMarkers() function. To account for the effect of ambient mRNAs, testing of ambient mRNAs was limited to cell types in which the ambient mRNA was a cell type marker gene.

#### Cell type resolved pathway analysis

The 1D annotation enrichment analysis^23^ was done with the freely available software package Perseus^57^, as previously described^28^. To predict the activity of upstream transcriptional regulators and growth factors based on the observed gene expression changes, we used the Ingenuity^®^ Pathway Analysis platform (IPA^®S^, QIAGEN Redwood City, www.qiagen.com/ingenuity) as previously described^28^. The analysis uses a suite of algorithms and tools embedded in IPA for inferring and scoring regulator networks upstream of gene-expression data based on a large-scale causal network derived from the Ingenuity Knowledge Base. The analytics tool ‘Upstream Regulator Analysis-™^22^ was used to compare the known effect (transcriptional activation or repression) of a transcriptional regulator on its target genes to the observed changes to assign an activation *Z-*score. Since it is *a priori* unknown which causal edges in the master network are applicable to the experimental context, the ‘Upstream Regulator Analysis’ tool uses a statistical approach to determine and score those regulators whose network connections to dataset genes as well as associated regulation directions are unlikely to occur in a random model^22^. In particular, the tool defines an overlap P-value measuring enrichment of network-regulated genes in the dataset, as well as an activation Z-score which can be used to find likely regulating molecules based on a statistically significant pattern match of up-and down-regulation, and also to predict the activation state (either activated or inhibited) of a putative regulator. In our analysis we considered genes with an overlap P-value of >7 (log10) that had an activation *Z*-score > 2 as activated and those with an activation *Z-* score < −2 as inhibited.

### Proteomics and multi-omics data integration

#### Detergent solubility profiling

For proteome analysis ~100mg of fresh frozen total tissue (wet weight) of mouse lungs was homogenized in 500 μl PBS (with protease inhibitor cocktail) using an Ultra-turrax homogenizer. After centrifugation the soluble proteins were collected and proteins were extracted from the insoluble pellet in 3 steps using buffers with increasing stringency as described in the QDSP protocol^28^. Peptides from LysC and trypsin proteolysis of the four protein fractions in guadinium hydrochloride (enzyme/protein ratio 1:50), were purified as previously described on SDB-RPS material stage-tips^28^.

#### Mass spectrometry

Data was acquired on a Quadrupole/Orbitrap type Mass Spectrometer (Q-Exactive, Thermo Scientific) as previously described^28^. Approximately 2 μg of peptides were separated in a four hour gradient on a 50-cm long (75-μm inner diameter) column packed in-house with ReproSil-Pur C18-AQ 1.9 μm resin (Dr. Maisch GmbH). Reverse-phase chromatography was performed with an EASY-nLC 1000 ultra-high pressure system (Thermo Fisher Scientific), which was coupled to a Q-Exactive Mass Spectrometer (Thermo Scientific). Peptides were loaded with buffer A (0.1% (v/v) formic acid) and eluted with a nonlinear 240-min gradient of 5–60% buffer B (0.1% (v/v) formic acid, 80% (v/v) acetonitrile) at a flow rate of 250 nl/min. After each gradient, the column was washed with 95% buffer B and re-equilibrated with buffer A. Column temperature was kept at 50 °C by an in-house designed oven with a Peltier element^58^ and operational parameters were monitored in real time by the SprayQc software^59^. MS data were acquired with a shotgun proteomics method, where in each cycle a full scan, providing an overview of the full complement of isotope patterns visible at that particular time point, is follow by up-to ten data-dependent MS/MS scans on the most abundant not yet sequenced isotopes (top10 method)^60^. Target value for the full scan MS spectra was 3 × 10^6^ charges in the 300-1,650 *m/z* range with a maximum injection time of 20 ms and a resolution of 70,000 at *m/z* 400. Isolation of precursors was performed with the quadrupole at window of 3 Th. Precursors were fragmented by higher-energy collisional dissociation (HCD) with normalized collision energy of 25 % (the appropriate energy is calculated using this percentage, and m/z and charge state of the precursor). MS/MS scans were acquired at a resolution of 17,500 at *m/z* 400 with an ion target value of 1 × 10^5,^ a maximum injection time of 120 ms, and fixed first mass of 100 Th. Repeat sequencing of peptides was minimized by excluding the selected peptide candidates for 40 seconds.

#### Mass spectrometry raw data processing

MS raw files were analyzed by the MaxQuant^61^ (version 1.4.3.20) and peak lists were searched against the human Uniprot FASTA database (version Nov 2016), and a common contaminants database (247 entries) by the Andromeda search engine^62^ as previously described^28^.

#### QDSP data analysis

The quantitative detergent solubility profiling (QDSP) analysis was done as previously described^28^. Briefly, intensities were first normalized such that the mean log2 intensities of the young and the old samples are zero, respectively. Using the normalized intensities, a two-way ANOVA with the two-factor treatment (old/young) and solubility fraction (FR1, FR2, FR3, INSOL) and the corresponding interaction term was performed using the R function aov(). Proteins significant in the interaction term correspond to proteins for which the solubility profile changes between young and old mice. Therefore, the corresponding P-value was used for filtering the significantly changed profiles after FDR correction.

#### *In silico* bulk differential expression analysis

In silico bulk samples were generated by summing UMI counts across all cells within one sample. Differential gene expression analysis of in silico bulk samples was performed using the R package DESeq2 (v1.20.0)^63^.

#### 2D annotation enrichment analysis

Statistical and bioinformatics operations, such as normalization, pattern recognition, cross-omics comparisons and multiple-hypothesis testing corrections, were performed with the Perseus software package^57^. The 2D annotation enrichment test used to compare proteome and transcriptome, is based on a two-dimensional generalization of the nonparametric two-sample test. The false discovery rate is stringently controlled by correcting for multiple hypothesis testing^23^.

### Flow cytometry

Isolated total lung cell suspensions were used to detect and quantify cell populations and activation by flow cytometry. We depleted red blood cells by positive selection of Ter199 cells, followed by CD45 bead separation (Miltenyi Biotec; Bergish Gladbach, Germany). Next, we analyzed cells by FACS cell suspensions before and after CD45 separation and stained cells suspensions with anti-mouse CD31, EpCAM, Podoplanin, (Biolegend; San Diego, CA, USA), H2-K1 (Thermo Fisher Scientific; Waltham, Massachusetts, USA). Cells were stained in the dark at 4°C, for 20 minutes. CD45 lineage negative cells were stained with Nile Red (Santacruz Biotechnology), in a 1:1000 dilution for 10 min at 4°C, as previously reported^64^. Data acquisition was performed in a BD Fortessa flow cytometer (Becton Dickinson; Heidelberg, Germany). Data were analyzed using the FlowJo software (TreeStart Inc; Ashland, OR, USA). Data were reported as absolute numbers (cells/uL), normalized by beads counts (BD Truecount TM Beads tubes; BD Biosciences, Heidelberg, Germany). For H2-K1 and Nile Red, data were analyzed by mean fluorescence intensity (MFI). Negative thresholds for gating were set according to isotype-labeled, and unstained controls.

### Immunofluorescence and histology

For immunofluorescence microscopy, mouse lungs were perfused with PBS, fixed in 4% paraformaldehyde (pH 7.0), and embedded in paraffin for FFPE sections. The paraffin sections (3.5 μm) were deparaffinized and rehydrated, and the antigen retrieval was accomplished by pressure-cooking (30 s at 125°C and 10 s at 90°C) in citrate buffer (10 mM, pH 6.0). After blocking for 1 h at room temperature with 5% BSA, the lung sections were incubated with the primary antibodies overnight at 4°C, incubated with the secondary antibodies (1:250) for 2 h, followed by DAPI (Sigma-Aldrich, 1:2,000) for 20 min at room temperature. Images were acquired with an LSM 710 microscope (Zeiss). The following primary (1) and secondary (2) antibodies were used: (1) CC10 rabbit (Santa Cruz, sc-25554), Foxj1 mouse (Santa Cruz, sc-53139), Collagen IV rabbit (Abcam, ab6586), Podoplanin goat (R&D Systems, AF3244); (2) donkey anti-mouse Alexa Fluor (AF) 647 (Invitrogen, A21447), donkey anti-rabbit AF 568 (Invitrogen, A10042), donkey anti-goat AF 488 (Invitrogen, A21202).

The frequency of ciliated (nuclear Foxj1+) and club cells (CC10+) were quantified by counting 2647 cells, covering a total length of 22 mm airway in 28 individual airways (young, n=14; old n=14) of two mice of each age group. We normalized cell numbers to the total length of their respective airway using the ZEN 2.3 SP1 software for image processing.

### Data availability

Proteome raw data can be downloaded from the PRIDE repository under the accession number XXX. scRNA-seq raw data can be downloaded from the Gene Expression Omnibus under the accession number XXX and accessed via the EBIs single cell expression atlas under the link XXX. The whole lung aging atlas can be accessed via an interactive user-friendly webtool at: https://theislab.github.io/LungAgingAtlas

## Acknowledgments

We thank Silvia Weidner and Daniela Dietel for excellent technical assistance. We also thank Gabi Sowa, Igor Paron and Korbinian Mayr for expert support of the proteomics pipeline. We thank Sandy Lösecke for technical assistance in next generation sequencing and Thomas Schwarzmayr for support with High-seq 4000 sequencing raw data. L.M.S. acknowledges funding from the European Union’s Horizon 2020 research and innovation programme under the Marie Sklodowska-Curie grant agreement No 753039. This work was supported by the German Center for Lung Research (DZL), the Helmholtz Association, and the German Federal Ministry of Education and Research (BMBF), project Single Cell Genomics Network Germany.

## Author contributions

H.B.S. conceptualized and supervised the entire project and wrote the paper. F.J.T. supervised single cell analysis and multi-omics data integration. I.A. and M.S. performed single cell transcriptomics experiments. L.M.S., I.A. and H.B.S. analyzed single cell transcriptomics data. H.B.S. performed proteomics experiments. H.B.S. and L.M.S. analyzed the proteomics data and performed transcriptomics and proteomics data integration. I.A. and C.H.M. performed histology and immunofluorescence microscopy. I.E.F. and F.R.G. performed flow cytometry experiments. E.G. and T.M.S. performed next generation sequencing of single cell libraries. L.M.S. and G.T. set up the interactive webtool. O.E. assisted in data interpretation and M.M. provided important support with mass spectrometry equipment. All authors read, edited and approved the manuscript.

---

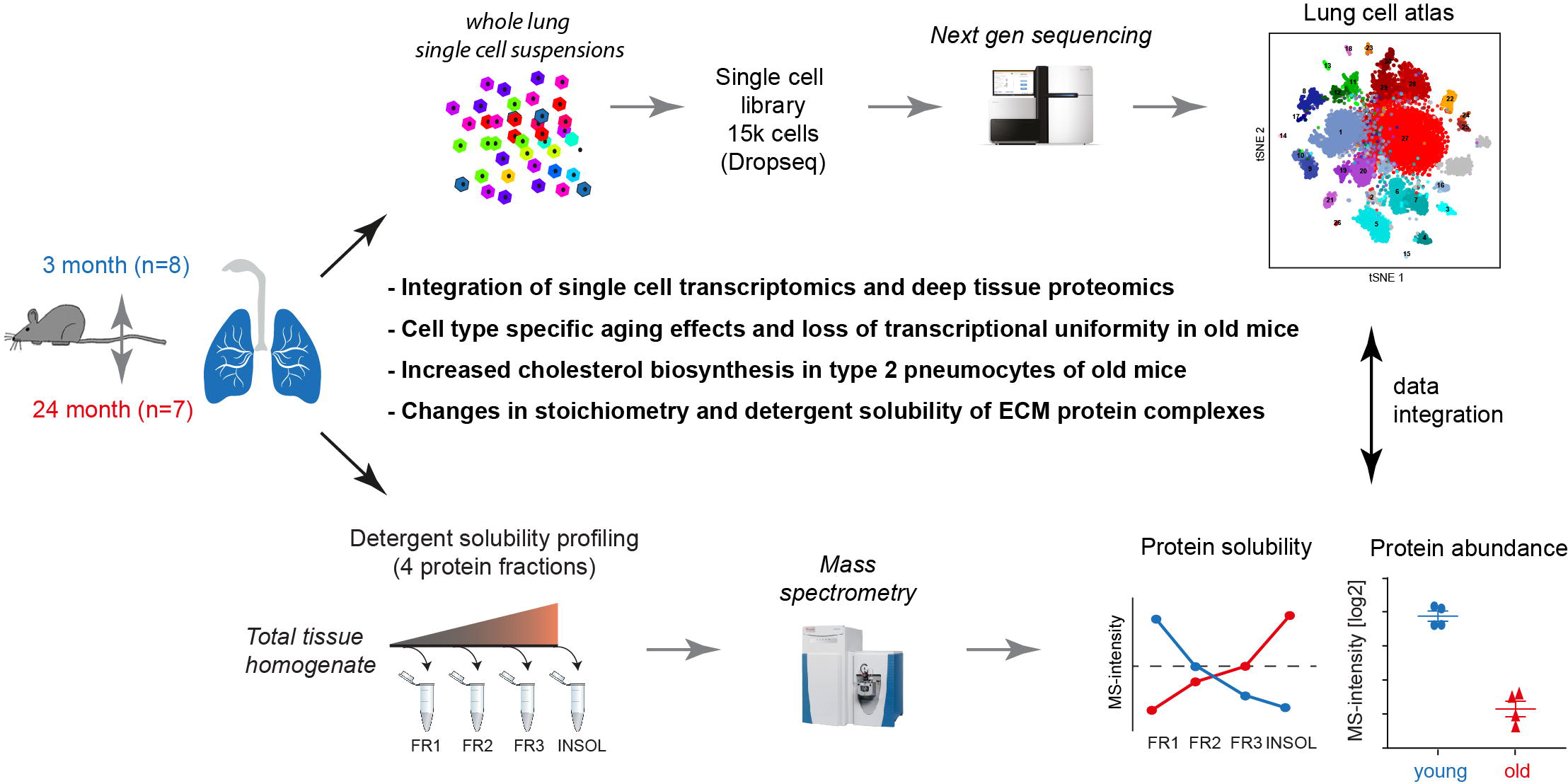

